# Inference of epistatic effects in a key mitochondrial protein

**DOI:** 10.1101/326215

**Authors:** Erik D. Nelson, Nick V. Grishin

## Abstract

We use Potts model inference to predict pair epistatic effects in a key mitochondrial protein – cytochrome c oxidase subunit 2 – for ray–finned fishes. We examine the effect of phylogenetic correlations on our predictions using a simple exact fitness model, and we find that, although epistatic effects are under–predicted, they maintain a roughly linear relationship to their true (model) values. After accounting for these corrections, epistatic effects in the protein are still relatively weak, leading to fitness valleys of depth 2*N*_*s*_ ~ −5 in compensatory double mutants. Positive epistasis is more pronounced than negative epistasis, and the strongest positive effects capture nearly all sites subject to positive selection in fishes, similar to virus proteins evolving under selection pressure in the context of drug therapy.

## I. INTRODUCTION

Epistasis in protein biophysics refers to the non–additive effects of amino acid mutations on a fitness trait [1], such as folding to a functional native state supporting a specific active site structure [2, 3]. For example, a deleterious mutation that disrupts folding may alter the statistical balance among conformation states available to a protein in such a way that a second, formerly deleterious mutation becomes beneficial in the new background [3], and the combined effect of the two mutations is nearly neutral – a form of positive, or compensatory epistasis [4]. Alternatively, mutations that are neutral, or nearly neutral individually may disrupt folding in combination – a form of negative epistasis. In either case, the combined effect of the mutations differs from the sum of their individual effects. This difference provides a measure of non–additivity which extends to any number of mutations [5].

Epistatic effects are thought to play a significant part in protein evolution [6–11]. However, because the conformational medium through which sites in a protein interact changes with each mutation, epistatic interactions between mutations are very difficult to predict [12]. In fact, even when all single and double mutation fitness effects in a protein are known, it may not be possible to predict the course of its evolution beyond a few mutations [13]. Explicit sampling of fitness and epistatic effects by mutagenesis is costly, and is limited to proteins that can be expressed and evolved in the laboratory. However, for certain proteins [14], the number of known sequences is sufficient to infer the fitness effects of mutations by machine learning methods – for example, by adjusting the parameters of a Potts spin model to recover the frequencies of amino acids in a protein alignment [14–17].

In the Potts inference approach, the probability of finding a particular sequence x in an alignment is assumed to follow a Boltzmann distribution,

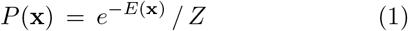

where

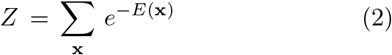

and

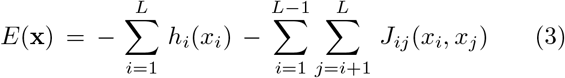

is the Potts energy of a sequence. Here, *x*_*i*_ is the amino acid state (i.e., type of amino acid) at site *i* in sequence x, and *L* is the length of sequences in the alignment (the sum in Eq. (2) runs over all possible amino acid sequences of length *L*). In order to fit the model to alignment data, the fields *h*_*i*_(*a*) and the couplings *J*_*ij*_(*a, b*) are adjusted until the one–point and two–point frequencies of amino acids states defined by the model agree with those of the alignment to within errors due to finite sampling (see below).

It can be shown that the equilibrium distribution of sequences generated by a Wright–Fisher process converges to a Boltzmann distribution when mutations are infrequent – specifically, when *μN* ≪ 1, where *N* is the effective population size and *μ* is the mutation rate per gene, per generation [18, 19]. In this limit, the time to fixation or loss of a mutation is much shorter than the time between fixation events so that fixation events usually take place against a monomorphic background. As a result, the evolution of a population can be described by the trajectory of a single representative sequence [20], similar to a Monte Carlo process [21]. The fitness of a sequence *f*(x) in the Wright-Fisher process is then related to its energy in the Boltzmann ensemble as (assuming *N* − 1 ≃ *N*)

**FIG. 1:**
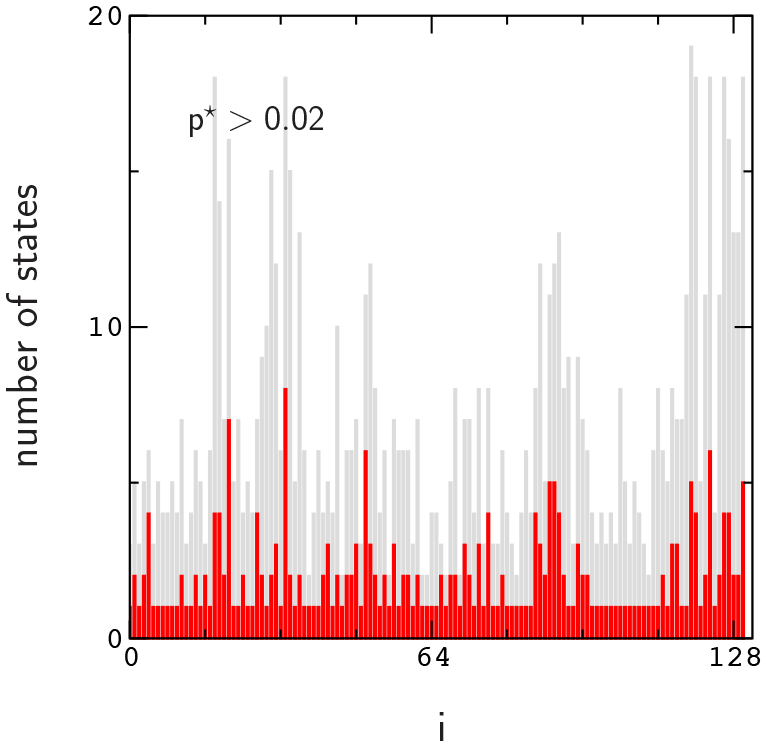
Number of observed states (grey), and number of states with 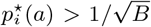 (red) for alignment (i). The predictive state space is defined by the set of sites with two or more states satisfying this inequality.

 
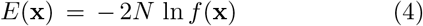

and energy differences between mutants x and x′ can be expressed as

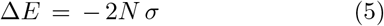

where *σ* = ln *f*(x′) − ln *f* (x). Here, the logarithmic fitness change *σ* is related to the selection coefficient, or relative fitness change *s* in the usual expression for the fixation probability obtained by Kimura [22] as *σ* = ln(1 + *s*). For realistic population sizes, N ≫ 1, and nearly neutral changes in fitness, *N*|*s*| ~ 1, the distinction between *σ* and *s* can be neglected, since ln(1 + *s*) ≃ *s* when |*s*| ≪ 1.

It is easy to see that these relations provide a rather tenuous connection to the forces which shape the distribution of real (naturally evolved) sequences in an alignment. Real sequences are not randomly sampled from an equilibrium distribution evolved under a constant fitness constraint *f*(x) at constant *μ* and *N* [19]. In a real phylogeny, sequences (leaf nodes of a phylogenetic tree) are correlated, and each of these quantities will have varied both along and among lineages due to genetic linkage, interactions among gene products, and changing environmental conditions [23–25]. As a result, fitness differences *σ* obtained from an inferred Potts model through Eq. (5) represent a kind of phylogenetic average over the effects of fluctuating constraints.

Recently, Shekhar et al. studied these effects using a model of HIV proteins evolving among human hosts [26]. In their model, proteins evolve according to an intrinsic fitness function of the form Eq. (3) plus a random field component representing host immune pressure that affects small groups of sites in a protein, randomly selected for each host. An encouraging result of their work is that the intrinsic coupling constants *J*_*ij*_(*a, b*) (which define pair epistasis in the Potts model) are recovered in a phylogenetic average over virus strains. Thus, we wondered whether the Potts inference method could be used to predict epistatic effects in a more general setting where sequences are sampled from a phylogeny of similar organisms.

**FIG. 2:**
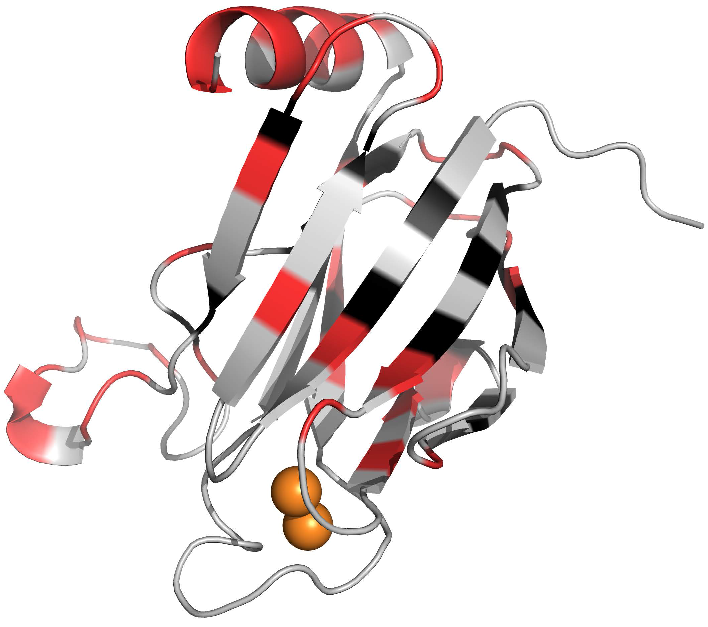
Predictive sites from Fig. 1 (red and black) mapped to the C–terminal domain of bovine heart COX2. Predictive sites with side–chains facing inward toward the core of the domain are colored black, otherwise they are colored red. Cu–atom ligands are shown as orange spheres.

Mitochondrial proteins provide a potentially interesting example [27, 28]. The evolution of mitochondrial proteins through different genetic backgrounds to some extent resembles the evolution of virus proteins in human hosts, and known sequences for mitochondrial proteins are plentiful. In this short article, we use the maximum entropy inference method developed for virus proteins [14] to predict pair epistatic effects in a mitochondrial protein – cytochrome c oxidase 2 – for ray-finned fishes.

Cytochrome c oxidase 2 (COX2) is the primary active subunit of cytochrome c oxidase (COX) – an enzyme consisting of 3 mitochondrial and 10 nuclear encoded proteins located in the mitochondrial inner membrane [29, 30]. Transient encounters between cytochrome c (CYC) and COX2 on the surface of the membrane transfer electrons to its interior, sustaining an electrochemical gradient that supports the production of ATP, which in turn supplies energy for processes such as muscle contraction and brain activity in fishes and other vertebrate organisms. Due to its role in energy production, it is thought that the evolution of organismal traits such as speed and endurance in high performance fishes may have been influenced by the adaptation of COX proteins [31, 32]. Accelerated rates of mutation indicating positive Darwinian selection have been observed at a number of sites in fish COX2, however, their involvement in COX activity is poorly understood. A number of these sites interact directly with nuclear subunits of COX, and most are distant from sites involved in the encounter with CYC. Mutation rates in mitochondrial proteins are also much larger than those for nuclear proteins leading to more frequent generation of deleterious mutations. These facts suggest a possible non–adaptive explanation for the acceleration of rates in terms of nuclear compensatory mutations [33]. However, because the effects of mutations are not always limited to the immediate vicinity where they occur in a folded protein (i.e., due to allostery, or long-range epistasis), the functional effects of the small bursts of mutations found in fish COX2 are essentially unknown.

**FIG. 3:**
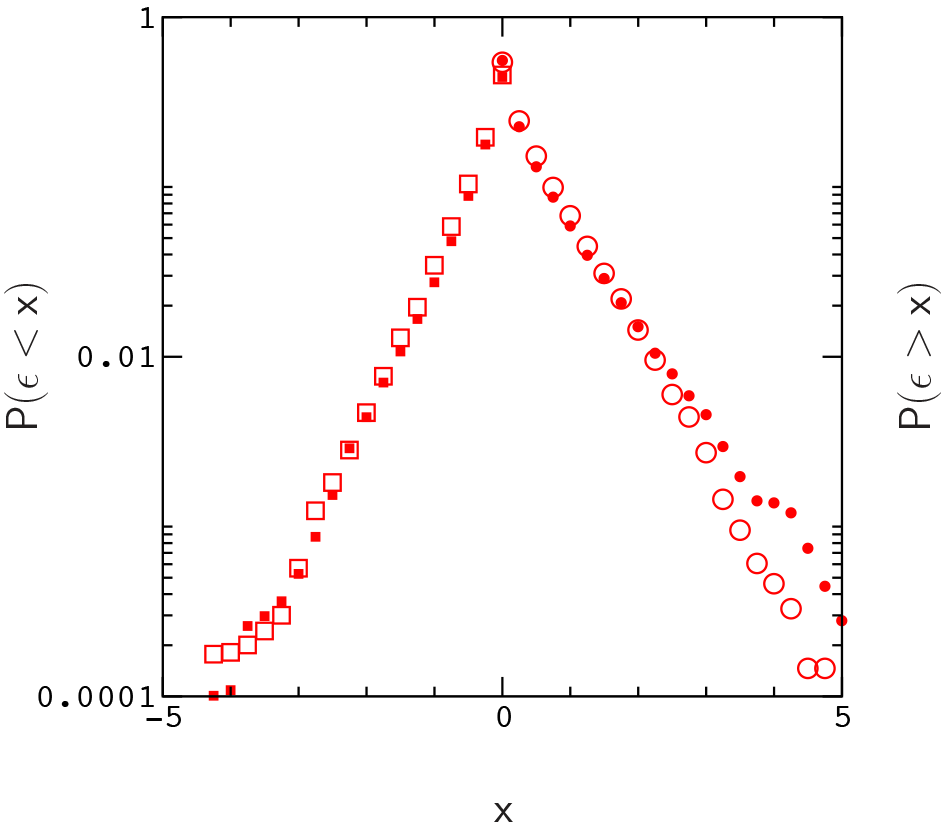
Frequencies *P*(*ϵ* > *x*) (circles) and *P*(*ϵ* < *x*) (squares) for positive and negative epistasis obtained for alignment (i) (open symbols) and alignment (ii) (filled symbols) with close homologs removed.

Existing data indicate that *μN* < 1 for mitochondrial genes in fishes, roughly consistent with the approximation in Eq. (5) (here, we have used N ~ 10^4^ for fishes [34–37], and pedigree rates *μ* ~ 10^−5^ for humans [38], which are more accurate, but probably a bit larger than those for fishes). These observations, together with those of Shekhar et al. [26] suggest that pair epistatic effects in fish COX2 sequences are predictable. Below, we compute the distribution of epistatic effects in fish COX2, and briefly explore the role of epistasis in COX2 evolution.

Let *F* denote the scaled logarithmic fitness of a sequence, *F* = 2*N* ln *f*, and let Δ*F* = 2*Nσ*. We define epistasis, using the expression in [5],

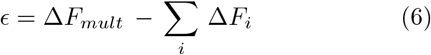

where Δ*F*_*i*_ and Δ*F*_*mult*_ are the individual and combined effects of mutations in a multiple mutation relative to a given “wild type” sequence (here we consider only pair mutations so that the sum in Eq. (6) contains only two terms, however, the notation is convenient for our discussion below). Epistasis is estimated from an inferred model by replacing Δ*F* with −Δ*E* in Eq. (6). We expect that weakly deleterious substitutions of strength Δ*F* ≃ −5 will occur fairly often in naturally evolved sequences [39, 40] (this can also be understood from the approximation in Eq. (5), assuming Δ*E* = −Δ*F* in the Metropolis acceptance condition [21]). Consequently, we expect to observe (for example) compensatory neutral epistatic effects of strength *ϵ* ≃ 10 in our inferred model (i.e., with Δ*F*_*mult*_ ≃ 0 and Δ*F*_i_ ≃ −5 in Eq. (6)) to the extent that such events were available in the ancestral states of proteins in our data. Instead, we find that, in general, predicted epistatic effects are limited to the range |*ϵ*| ⪝ 5.

**FIG. 4:**
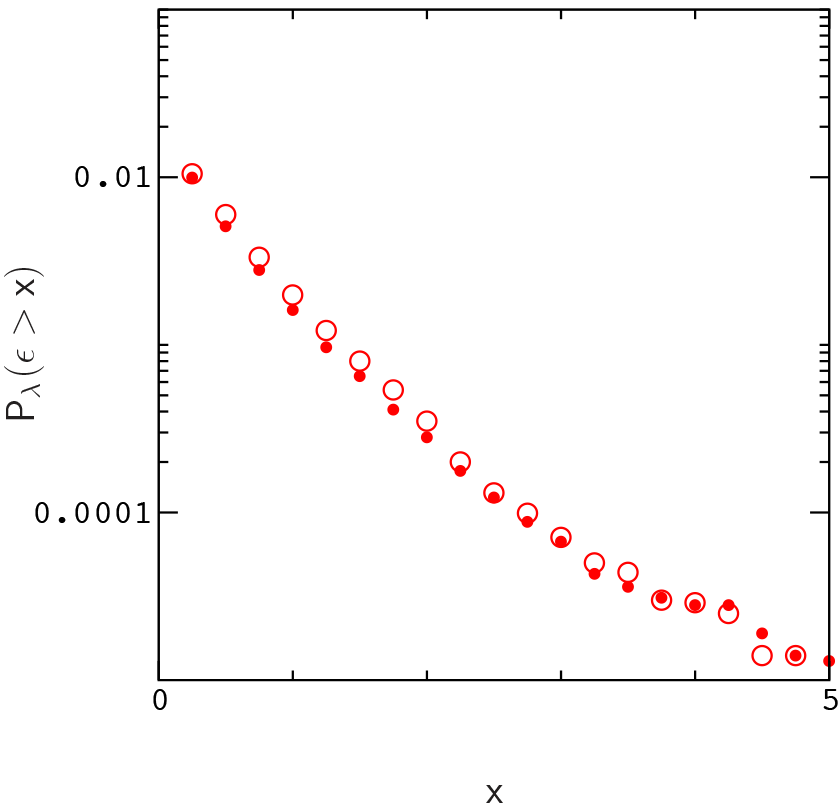
Frequencies *P*_λ_(*ϵ* > *x*) for compensatory, nearly neutral fitness epistasis with Δ*F*_*mult*_ > −λ (λ = 2) for alignments (i) (open circles) and (ii) (filled circles).

To see how phylogenetic correlations and limited (”non-equilibrium”) sampling of the fitness landscape could have affected our predictions, we evolve a simple exact fitness model of the form Eq. (3) to reflect the structures of phylogenetic trees derived from sequences in our input alignments. The results of these simulations suggest that epistasis in COX2 is under-predicted by a factor of order 2. Hence, after including this correction, epistatic effects with |*ϵ*| ≃ 10 occur at a low frequency. Interestingly, positive epistasis is more prevalent than negative epistasis, and the strongest positive effects capture nearly all nominally adaptive sites in fishes, similar to virus proteins evolving under selection pressure in the context of drug therapy [6–8], which suggests that the asymmetry in epistasis may have adaptive origins.

We map the strongest positive epistatic interactions onto the crystal structure of COX2 and find that they often connect sites with accelerated mutation rates to segments of the protein that are more directly involved in the encounter with CYC, however, our results are sensitive to the choice of sequences included in alignments.

In the following sections, we first discuss the inference method in more detail and describe the procedure used to process our input sequence data. In subsequent sections, we present and briefly discuss our results.

## II. METHODS

To obtain an inferred fitness model, fields, *h*_*i*_(*a*), and couplings, *J*_*ij*_(*a, b*), are adjusted until the frequencies

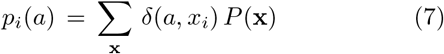

and

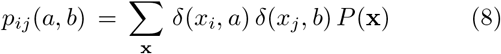

of amino acid states described by the model agree with the frequencies

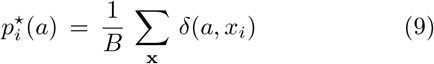

and

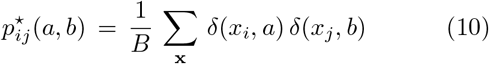

and we choose *q* so that *S*_*q*_ ≥ 0.9 *S*, where *S* is the total entropy of the site. The first *q* states are treated explicitly, while the remaining states (excluding the least probable state) are mapped to a single compressed state, *σ*_*i*_ – the least probable state being reserved for what is known as the “gauge state” in the inference method (a more detailed expanation of this procedure can be found in ref. [14]). Frequencies are re-computed to reflect the mapping, and the fields and couplings are inferred of amino acid states in a multiple sequence alignment (MSA). Here, *B* is the number of sequences in the MSA, *δ*(*a, b*) = 1 if states *a* and *b* are the same, and *δ*(*a, b*) = 0 otherwise. To solve the inference problem, we use the adaptive cluster expansion (ACE) and Boltzmann machine learning algorithms developed by Barton et al., which avoid over-fitting by constructing a sparse network of interactions sufficient to reproduce the observed frequencies to within errors due to finite sampling [15] (see Jacquin et al. [16] for a comparison different methods).

The threshold for prediction of the coupling constants *J*_*ij*_(*a, b*) is roughly determined by the condition 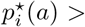 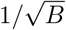 [15]. However, even for large samples, the parameter space {*J*_*ij*_(*a, b*)} may be too complex to obtain complete model including all amino acid states sampled in the MSA. In this case, which is the norm for proteins, a natural way to reduce the size of the parameter space is to group infrequently observed states in an alignment column – typically with 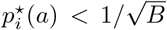 – into a single “compressed” state.

Following Barton et al. [14], we group infrequent states so as to retain 90 percent of the site entropy. For each site, amino acid states *a*_*i*_ (including a gap state) are first arranged in order of decreasing frequency,

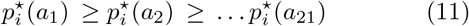

The Shannon entropy of the first *q* states plus the compressed state can then be expressed as,

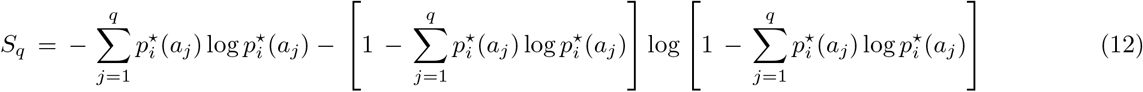

from the re-computed frequencies. Parameters involving states *a* ∈ {*σ*_*i*_} are then re-constructed from the inferred parameters as 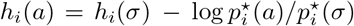 and *J*_*ij*_(*a, b*) = *J*_*ij*_(*σ, b*), etc., where *σ* is the compressed state for site *i*.

To obtain our input MSA, we used BLAST to interrogate the UniProtKB database against COX2 from sailfish (Istophorus albicans), a member of a recently diverged branch on the phylogenetic tree for fishes [41].

**FIG. 5:**
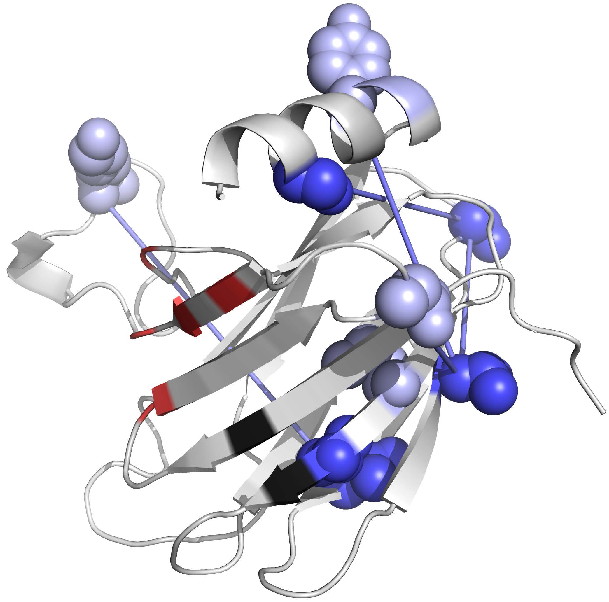
Epistatic interactions with *ϵ* > 3.5 involving sites with accelerated mutation rates (dark blue) for the model obtained from alignment (i). Interacting sites are linked by bonds. Sites without accelerated rates are colored light blue, and sites that interact directly with CYC in reference [29] are colored red. The most frequently observed epistatic pair is indicated in black (these sites do not have accelerated rates). Identical results are obtained for *ϵ* > 3.5 and Δ*F*_*mult*_ > −2.

This choice has the effect of segregating fish sequences from those of other organisms, and for convenience, we simply selected the 3600 best hits for subsequent alignment, including a small fraction of amphibian and tetrapod sequences. Sequences were aligned using ProMALS-3D [42]. Alignment columns containing more than fifty percent gaps were removed, and unknown residue types were replaced by entries from corresponding sites in the closest available sequence. This last step affects a small number of sequences, usually at the C–terminal end of the protein. Gaps are retained as possible states of the Potts model, and are, likewise, usually found at or near the C–terminal end. To minimize bias introduced by repeated sampling of certain organisms (e.g. salmon, cod, etc.) we selected a single sequence from each group of identical sequences in the alignment. This step results in about 2600 unique sequences.

Below, we model only the globular domain region of COX2, neglecting the N-terminal helices (see Fig. 2), and we refer to the alignment above (with N–terminal segments removed) as alignment (i). We then repeat the step above, selecting a single sequence from each group of identical sequences in alignment (i), in effect removing close homologs. This step results in about 2300 unique sequences that we refer to as alignment (ii). Below, we compute epistatic effects for both alignments in order to test the robustness of our predictions. Fields and couplings are inferred by straight forward application of the adaptive cluster expansion and Boltzmann machine learning algorithms using weak regularization (γ ≃ 2/*B*) similar to the approach used for proteins in ref. [15].

**FIG. 6:**
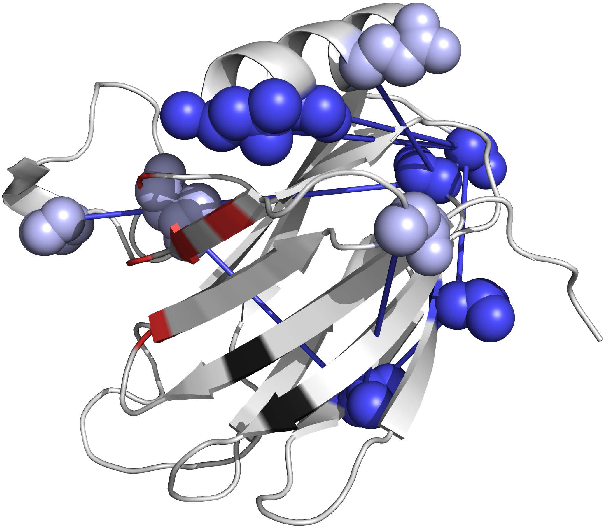
Epistatic interactions with *ϵ* > 3.5 for the model obtained from alignment (ii). The layout of the plot is the same as in Fig. 5. Identical results are obtained for *ϵ* > 3.5 and Δ*F*_*mult*_ > −2.

## III. RESULTS

To survey the effects of pair mutations, we restrict amino acid transitions to states in the predictive subset 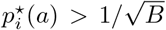, and therefore, to sites that have at least two states satisfying this condition (Fig. 1). In Fig. 2, we map these “predictive sites” onto the structure for bovine heart COX2 by aligning our MSA sequence for sailfish to the sequence obtained from the crystal record (Protein Data Bank id 3abk). Predictive sites indicated in Fig. 2 (colored red and black) include about 20 amino acids that face inward toward the core of the *β*–domain (black), and among these sites there are about 40 instances of close approach between amino acids in which side-chain surfaces are separated by less than an atomic diameter. Many more instances of close approach, or contact occur between amino acids within loops external to the core, and on the surface of the *β*–domain (red), hence, the connectivity of this set presents many opportunities for short-range compensatory epistasis [4, 40]. By contrast, sites excluded from the predictive set are usually located within the binding interfaces joining COX2 to other COX subunits (specifically, subunits 1 and 6B1) or in loops attached to the metal ligands (orange spheres in Fig. 2). A similar division of sites into mutable and conserved groups is obtained in terms of the site entropy (see Supplemental Material [43]).

In Fig. 3, we plot the distributions *P*(*ϵ* > *x*) and *P*(*ϵ* < *x*) for positive and negative epistasis derived from alignments (i) and (ii). Note that positive epistasis is more pronounced when close homologs are removed. For comparison (Fig. 4), we plot the distribution for positive epistasis with double mutant fitness effects Δ*F*_*mult*_ restricted to Δ*F*_*mult*_ > −λ with λ = 2 – i.e., the frequency of compensatory nearly neutral, or beneficial double mutants, *P*_λ_ (*ϵ* > *x*). To construct the figures, we sample all sequences in the initial alignment – explicitly, for each sequence, we sample all distinct pairs of amino acid mutations in the predictive space, 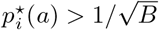, restricted to those allowed by the genetic code for vertebrate mitochondria (i.e., in a transition from amino acid state *a* to state *b*, at least one codon for *b* must be accessible from a codon for state *a* by a single base change). Nearly identical plots are obtained using much smaller, random samples of sequences.

**FIG. 7:**
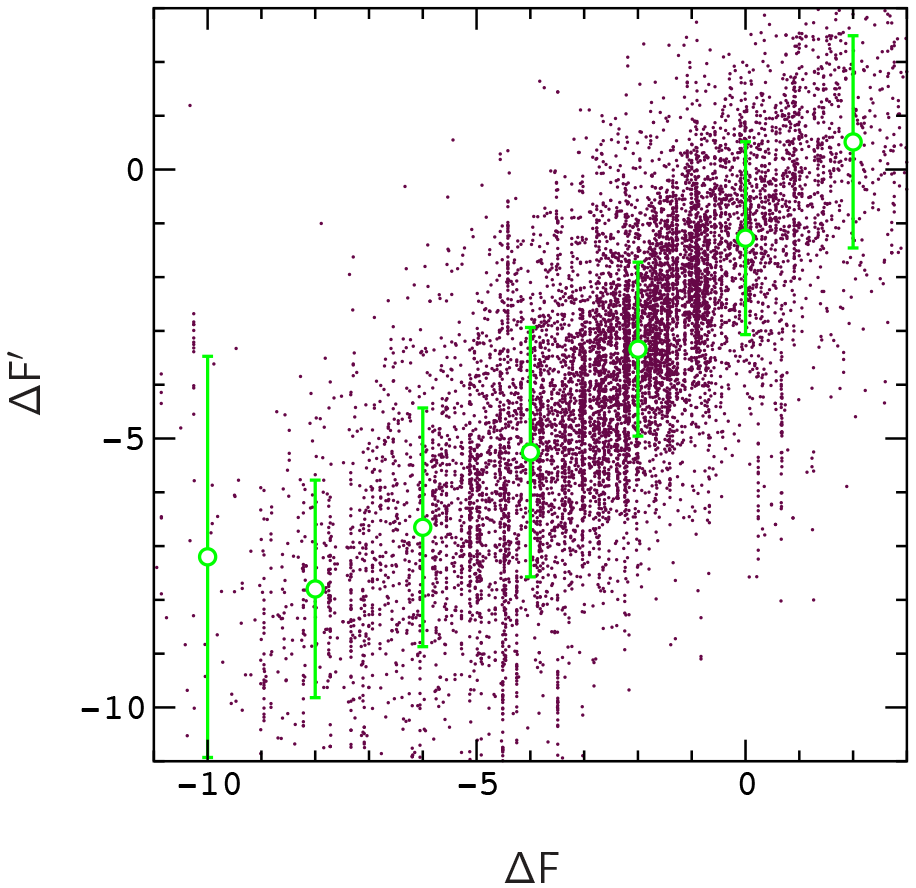
Comparison of predicted (Δ*F*′) and actual (Δ*F*) fitness changes obtained by tracing the tree for alignment (i). Circles and error bars indicate the average and standard deviation of Δ*F*′ for a given value of Δ*F*. Standard errors fit within the symbols (circles) used to indicate the averages, and are omitted for clarity. The scattered data is a random sample of the points used to compute the averages.

To explore the potential role of positive epistatic interactions in COX activity, we project the strongest effects, *ϵ* > 3.5, onto the bovine crystal structure. This condition captures 6 out of 7 sites in the C–terminal domain with accelerated mutation rates in each inferred model [31, 32]. Interactions that involve these sites are shown for the model obtained from alignment (i) in Fig. 5, and for alignment (ii) in Fig. 6. Sites involved in epistatic interactions are linked by bonds, and sites with accelerated mutation rates are colored dark blue (sites that interact directly with cytochrome c are colored red). In general, each bond represents a number of different pair mutations and organismal sequences. For both models, bonds tend to connect sites with accelerated rates to segments of the protein that border the encounter region (e.g., the *β*–hairpin containing the encounter sites, and the C-terminal helix, which interacts with the hairpin). However, the pattern of bonds changes when close homologs are removed (Fig. 6). The most frequently observed interaction in both models involves a contact between sites 100 and 155 on the binding interface (black) that do not have accelerated mutation rates.

**FIG. 8:**
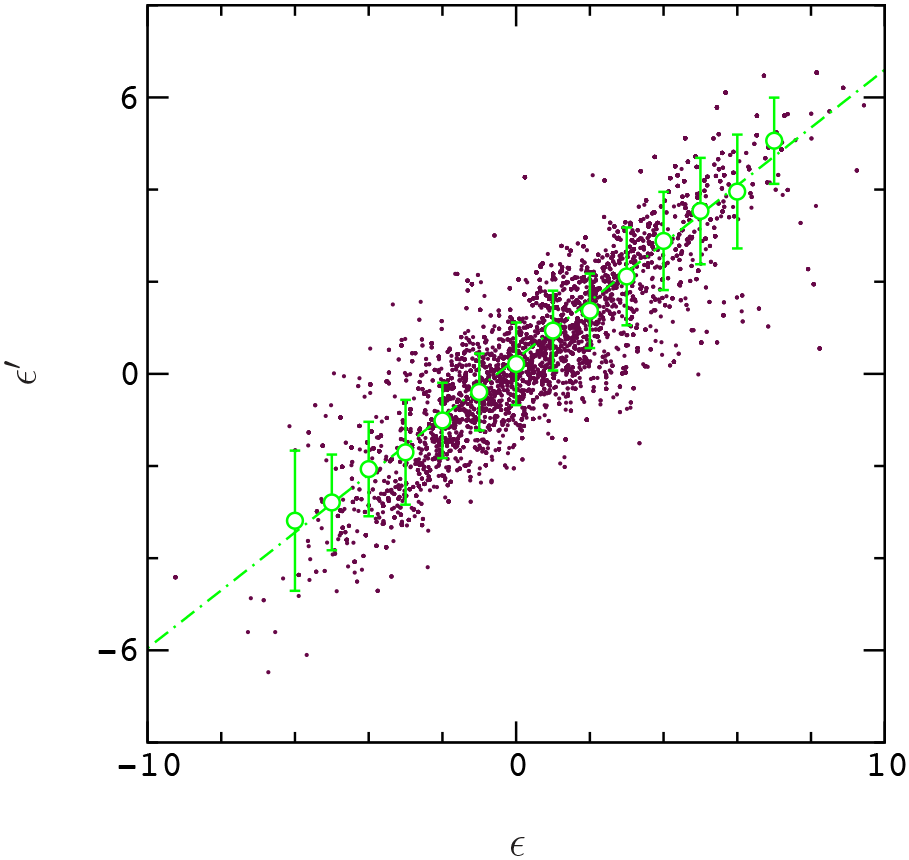
Comparison of predicted (*ϵ*′) and actual (*ϵ*) values of epistasis obtained by tracing the tree for alignment (i). The layout of the plot is the same as in Fig. 7. Standard errors fit within the symbols (circles) used to indicate the averages, and are omitted for clarity. The dashed line is a linear fit to the averages.

To estimate the effects of phylogenetic correlations and limited sampling on the scale of epistatic effects in our inferred models, we evolve sequences to reflect the topologies of binary phylogenetic trees obtained from alignments (i) and (ii) using a simple exact fitness model of the form, *F*(x) = −*αE*(x), where *E*(x) has the form Eq. (3) (more details are provided in the Appendix).

To initiate this process, a tree is constructed for alignment (i) or (ii), and a single well equilibrated model sequence is created to represent the root node of the model tree. At the first branching node of the real tree, the sequence is split into two copies, which are then evolved for periods of time proportional to the emergent branch lengths. This procedure is continued until all branches of the tree are “traced” to their leaf nodes, and the resulting ensemble of sequences is used to predict the model fitness function *F*(x) (here the procedure used for inference is the same as above). For clarity, we construct trees for alignments (i) and (ii) using the UPGMA method, and for simplicity, sequences are evolved by Monte Carlo (for the validity of this approximation, see the Appendix and ref. [21]). The energy scale of the model, *α,* is set to obtain a biologically meaningful acceptance rate (≃ 0.05), and the constant of proportionality between branch length and sampling time, *β*, is set to obtain a similar value for the mean distance (per mutable site) for real and traced sequences. The tracing process does not, of course, re-capitulate the structures of the input trees, however, the model trees generated by tracing have broad distributions of branch lengths, similar to real trees (expanded diagrams for real and model trees are included in the Supplementary Material [43]).

Predicted and actual values of Δ*F* and *ϵ* obtained by tracing the tree for alignment (i) are compared in Figs. 7 and 8. Predicted values of *ϵ* maintain a roughly linear relationship to their true values, however, *ϵ* is underpredicted by a factor of order 2 (very similar plots are obtained by tracing the tree for alignment (ii)). Results for equilibrium sampling, and for smaller values of the proportionality factor *β* are described in the Appendix.

To conclude, after accounting for under-prediction due to phylogenetic correlations and limited sampling of the fitness landscape (Fig. 8), epistatic effects in the inferred models are consistent with what one might expect from evolution of a protein by Wright-Fisher dynamics in the *μN* ≪ 1 limit. Positive epistasis is more prevalent than negative epistasis (Fig. 3), and the strongest positive effects tend to be associated with nominally adaptive sites identified in [31, 32]. A similar situation occurs for certain virus proteins evolving under strong selection pressure [6–8], which suggests that the asymmetry in Fig. 3 could be a residual effect of early adaptive events in fishes. The results for specific epistatic interactions in Figs. 5 and 6 seem to support a connection between accelerated mutation rates in COX2 and CYC binding efficiency (i.e., adaptation), but the evidence for this is relatively weak. A more thorough study of these problems, including selection pressure in a realistic fitness model such as that in [13], could be very useful to understand the dynamics of epistasis in protein populations, and the sources of error in model inference of epistatic effects.

## Acknowledgements

It is a pleasure to thank John Barton for many helpful conversations on the inference method, Wenlin Li and Jimin Pei for help with phylogeny methods, Fyodor Kondrashov for sharing his sequence data from ref. [27], and one anonymous reviewer for a number of helpful suggestions. This work was supported in part by the National Institutes of Health (GM094575 to NVG) and the Welch Foundation (I-1505 to NVG).

## Appendix

To estimate the effect of phylogenetic correlations and limited sampling of the fitness landscape on the scale of epistatic effects in our inferred models, we trace the phylogenetic trees obtained from alignments (i) and (ii) using a random network fitness model, *F*(x) = −*αE*(x). Here *E*(x) is given by the 20–state Erdos-Renyi energy model studied in [15] and the factor α ≃ 1.3 is included to obtain a biologically meaningful acceptance rate (≃ 0.05) in the tracing procedure. Fields *h*_*i*_(*a*) and couplings *J*_*ij*_(*a, b*) in the model are randomly selected from gaussian distributions of width *σ*_*h*_ = 1 and 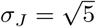. Sites in the network interact if they are connected by an edge, and edges are included at random with probability *p* = 0.05 (the model has *L* = 50 sites).

UPGMA trees for alignments (i) and (ii) are generated using the neighbor command in the *phylip* software library [44], and trees are traced by Monte Carlo (i.e., assuming an exponent −*αE*(x) in the Boltzmann factor). Distributions *P*(*d* > *x*) of Hamming distances d(x, x′) between leaf node sequences of real and model trees obtained for alignment (i) are compared in Fig. 9 (tree diagrams are included for comparison in the Supplementary Material [43]). Although the tracing procedure (see text) generates broad distributions of branch lengths, the branches of model trees are more homogeneous (i.e., long (short) branches in “real” trees tend to be slightly shorter (longer) than those in model trees). Smaller constants of proportionality between branch length and sampling time *β* lead to more homogeneous branch lengths (Fig. 9), and smaller predicted values for *ϵ*, however, the roughly linear relationship between *ϵ*′and *ϵ* exhibited in Fig. 8 is maintained. It is important to remark that, while the Monte-Carlo process is equivalent to the *μN* ≪ 1 limit of the Wright-Fisher process in equilibrium, it tends to accept smaller changes in fitness (energy) at a slightly higher rate, which may lead to smaller inferred values of Δ*F* and *ϵ* (for a comparison, of these processes, see ref. [21]). Finally, for sequences sampled from an equilibrium distribution, *ϵ* is still under-predicted, with *ϵ*′/*ϵ* typically of order 8/10 (this may to some extent reflect the regularization penalty in the inference method [15], however, we did not explore this issue in detail). For equilibrium sampling, sample sizes ranging from 3k to 10k sequences produced qualitatively similar results.

**FIG. 9:**
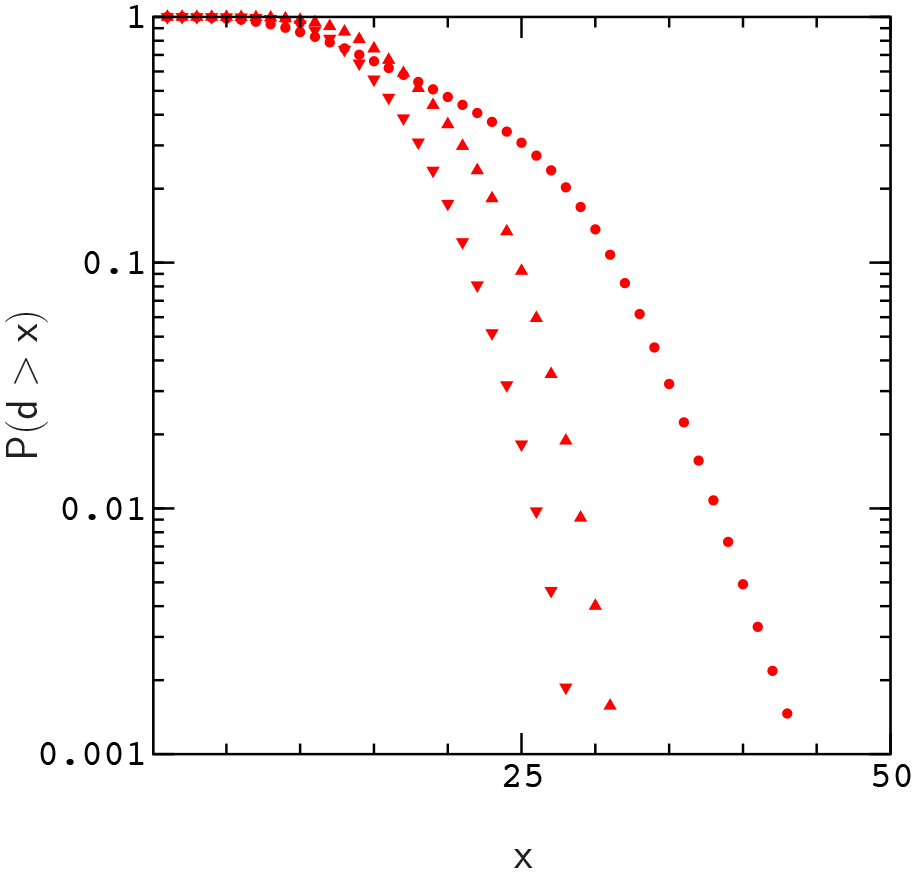
Distribution *P*(*d* > *x*) of Hamming distances, *d*(x,x′), for sequences in alignment (i) (circles), model sequences obtained by tracing the tree for alignment (i) in Figs. 7 and 8 (upward triangles), and model sequences obtained by tracing the tree for alignment (i) with a smaller value of *β* (downward triangles).

